# Quantitative Gait Analysis in a Sheep Model of In-Utero Spina Bifida Repair

**DOI:** 10.64898/2026.09.16.752110

**Authors:** Tsung-Yeh Chou, Shreya Pandiri, Payam Zandiyeh, Luis A. Figueroa Fernandez, Dejian Lai, Ramesha Papanna, Lovepreet K. Mann

**Author notes:** **Corresponding Author:** Lovepreet K. Mann, MBBS, Department of Obstetrics, Gynecology & Reproductive Sciences, McGovern Medical School at UTHealth Houston, 6431 Fannin Street | MSB 3.262 **|** Houston, Texas 77030, tel **|**.

## Abstract

**Objective:** To evaluate quantitative gait analysis in a sheep model (*Ovis aries*) of in-utero spina bifida repair.

**Method:** Using a 3D marker-based motion capture system, we quantified spatiotemporal gait parameters and hindlimb joint angles in a sheep model of in-utero spina bifida repair and normal controls. A linear support vector machine classifier, trained and tested on subject-disjoint partitions with individual gait cycles as the unit of analysis, evaluated the relationship between hindlimb joint angles and clinician-assigned Texas Spinal Cord Injury Scale scores. Linear mixed-effects models compared spatiotemporal parameters, hoof height, and hip drop across 3- to 6-month evaluations and between groups.

**Results:** In treated (n=14) and control (n=3) lambs, front- and hindlimb gait speed differed by timepoint (both p=0.025). Across 1,632 left and 1,661 right hindlimb gait cycles, held-out classification accuracy was 85.9% and 88.6% (Cohen’s κ=0.78 and 0.84) for left and right hindlimbs, respectively, against majority-class proportions of 76.5% and 75%. Left hindlimb hip drop differed between treated lambs and controls (Δ=0.8 cm, p=0.027). Between-group contrasts were considered exploratory given the small control group.

**Conclusion:** Quantitative gait analysis was feasible in an in-utero spina bifida repair model. Hindlimb kinematic profiles classified clinical ambulation scores above the majority-class baseline.

**What’s already known about this topic?:**

- Independent walking is a major benefit of in-utero spina bifida repair surgery and a driver of treatment innovation.
- Ambulation after in-utero repair is typically assessed using subjective clinical assessments, which limits the ability to evaluate the effect of treatment.
- Instrumented gait analysis enables precise evaluation of ambulatory function in three-dimensional movements but has not yet been implemented in models of in-utero spina bifida repair.

**What does this study add?:**

- We objectively quantified spatiotemporal gait parameters and hindlimb kinematic profiles over serial assessments in an established sheep model of in-utero spina bifida repair using a three-dimensional motion capture system.
- Hindlimb kinematic profiles classified clinician-assigned Texas Spinal Cord Injury Scale scores with 85.9%–88.6% accuracy, above the 75%–76.5% majority-class baseline. Pastern kinematics contributed most to classification.
- Treated lambs’ gait deviated from controls mainly during swing and mid-stance phases (exploratory).

## Introduction

The randomized Management of Myelomeningocele Study helped establish in-utero repair as the standard of care for spina bifida, in part by demonstrating improved neurological and functional outcomes after prenatal compared to postnatal repair.^1,2^ More children could walk independently at 30 months after prenatal than postnatal repair (44.8% vs. 23.9%, respectively),^3^ a benefit that persisted into school age.^4,5^ However, more than half of children in the prenatal repair group remain unable to walk independently.^1,3^ This persistent deficit has motivated the development of novel strategies to improve ambulatory function after in-utero repair,^6–13^ including our work investigating cryopreserved human umbilical cord as a meningeal patch.^14–18^

Nonetheless, these interventions’ effect on ambulatory function is typically evaluated using subjective clinical assessments. In the sheep model (*Ovis aries*) of in-utero spina bifida repair, the principal large-animal model for developing and testing fetal repair strategies,^19–23^ postnatal ambulation is mostly assessed using categorical scales.^15–17,24–27^ These assessments, which rely on subjective examiner judgment, lack the resolution to detect subtle gait abnormalities or to track their evolution over time, thus limiting the ability to evaluate treatment effect.

Instrumented gait analysis enables precise evaluation of ambulatory function in three dimensions throughout the gait cycle,^28^ providing sensitivity to detect subtle abnormalities involving multiple levels, joints, and anatomic structures that clinical observation may not capture. Detecting these subtleties in ambulation after in-utero spina bifida repair could guide the development and optimization of treatments. However, although 3D gait analysis has been applied in sheep models of spinal cord injury,^29–32^ these capabilities have not been applied to the sheep model of in-utero repair.

Here, we implemented a marker-based 3D motion capture system with machine learning analysis to objectively quantify gait in a sheep model of in-utero spina bifida repair. We assessed the feasibility of serial quantitative gait measurements in this model and evaluated the agreement between objective kinematic profiles and clinical scoring via the Texas Spinal Cord Injury Scale.

## Methods

### Study design and approvals

To evaluate the feasibility of objective serial gait assessments after in-utero spina bifida repair, we conducted a secondary analysis of a study comparing in-utero spina bifida repair methods in a sheep model. The UTHealth Houston Institutional Animal Care and Use Committee approved the study protocols (AWC-20-0149, approved May 11, 2021; AWC-23-0115, approved January 10, 2024). Animal care was conducted in compliance with the Guide for the Care and Use of Laboratory Animals.

### Sheep model of in-utero spina bifida repair

Timed-pregnant sheep with ultrasound-verified twin or triplet gestations underwent surgery at gestational day 75 (of 145) to create fetal lamb models of spina bifida without myelotomy, as described previously.^17,19,33^ The procedure was repeated on the fetus(es) in the remaining uterine horn(s) before skin and fascial incisions were closed. Fetuses that survived to repair were randomly assigned to one of three methods of in-utero repair, performed at gestational day 96: conventional repair (dura-myofascial closure) or repair using bovine collagen matrix (Durepair™, Medtronic, Minneapolis, MN) or cryopreserved human umbilical cord (TissueTech™, Inc., Miami, FL) allografts as a meningeal patch before skin closure, as described previously.^17^ Fetuses that did not undergo spina bifida creation served as controls. Lambs were delivered by vaginal or cesarean delivery on gestational day 140. Neonates were transitioned to room air with routine care, then stimulated for ambulation.

### Acclimation for gait protocol

Beginning four weeks after delivery, ambulatory sheep were acclimated to the gait protocol via daily training. Research staff guided sheep on a leash along a 10-meter mat walkway in the laboratory to familiarize them with the testing environment and protocol. Once sheep were comfortable with the environment and could walk continuously along the walkway for 10 minutes, training frequency was reduced to twice per week.

### Gait assessment via TSCIS

Once per month, two research members blinded to group allocation independently assessed animals’ hindlimb gait using the Texas Spinal Cord Injury Scale (TSCIS) during a 5-minute data collection period.^15–17,27^ The TSCIS evaluates each limb for gait, proprioception, and nociception (pain response). We scored only the gait subscale for comparison with quantitative gait parameters, and only the hind limbs as relevant to human ambulation. Gait was scored on a 0-6 scale, where 0 denotes no voluntary movement and 6 denotes normal gait. Scores from both raters were averaged.

### Motion capture data collection

Motion capture assessments were collected monthly, from 3-6 months after birth, using a 10-camera motion capture system (Qualisys, Gothenburg, Sweden; 100 Hz). Before each assessment, animals were shaved to ensure secure marker placement. Twenty-three reflective markers were placed on anatomical landmarks: five markers along each limb, laterally to the center of each joint, and one marker each on the head, thoracic, and sacral regions (Figure 1). Markers were always placed by the same research staff member to control inter-rater variability in landmark identification. Sheep completed a 5-minute familiarization trial to adjust to the handler, markers, and testing environment. Then, sheep walked for another 5 minutes for the testing trial.

**Figure 1.**
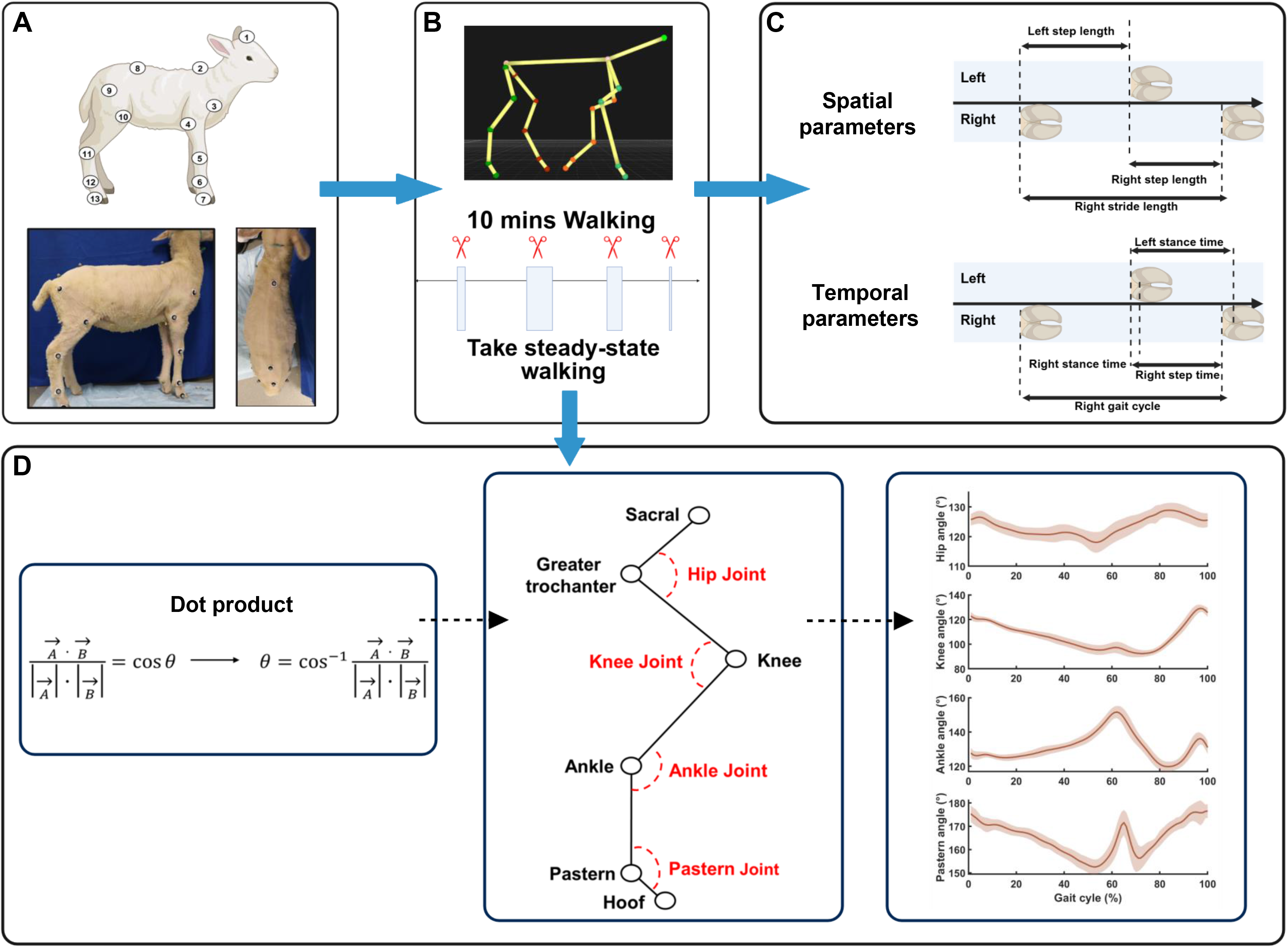
Analysis pipeline for spatiotemporal gait parameters and hindlimb sagittal-plane joint angles. (A) Schematic diagram shows the locations of reflective markers: 1) head; 2) thoracic spine; 3) glenoid; 4) elbow; 5) wrist; 6) sesamoid; 7) front hoof; 8) sacral region; 9) greater trochanter; 10) knee; 11) ankle; 12) pastern; and 13) hind hoof. (B) Steady-state walking blocks were selected from 10-minute walking trials. (C) Spatiotemporal gait parameters were calculated from selected walking data. (D) Hindlimb sagittal plane joint angles were calculated using a dot-product approach between adjacent segment vectors to quantify pastern, ankle, knee, and hip joint kinematics across the gait cycle.

### Data reduction and gait analysis

Motion capture data were screened to include only steady-state walkway passes with consecutive, uninterrupted steps, excluding the first and last 2 m of the 10-m walkway. Gait events were manually identified by the same research staff member using pastern markers. Initial ground contact was defined as the frame corresponding to the lowest vertical position of the pastern marker; toe-off was identified when the pastern marker began to rise from this position. The gait cycle was defined as the interval between consecutive heel strikes, identified using the pastern markers. A minimum of six steady-state gait cycles per limb per session were included for analysis. This threshold was derived via Generalizability Theory analysis of the number of gait cycles needed to obtain reliable gait outcomes (Supplementary Material 1).^34^

Marker trajectories were low-pass filtered with a zero-lag, fourth-order Butterworth filter (8 Hz cutoff frequency). The filtered marker positions were used to define segment vectors and compute sagittal-plane joint angles for the hip, knee, ankle, and pastern (Figure 1). Joint angles were calculated using vector dot-product methods to determine the angle between adjacent segments. Data were processed in Visual3D (HAS-Motion, Ontario, Canada) by an analyst blinded to group allocation and TSCIS scores.

Quantitative gait outcomes included spatial (step length, step width, and hoof height) and temporal (cycle time, step time, stance time, and swing time) gait parameters and joint angles for each limb and hip drop for hind limbs, for each assessment time point (3, 4, 5, and 6 months). Hoof height was defined as the maximum vertical position of the hoof marker during the gait cycle. Hip drop was defined as the vertical displacement between the ipsilateral sacral and greater trochanter markers during the stance phase of the contralateral limb. Positive values indicate a downward displacement of the pelvis on the ipsilateral side (hip drop), whereas negative values indicate an upward displacement (hip hike). Step length, step width, and hoof height were normalized to limb length. The reliability of each gait outcome was evaluated using Generalizability Theory to quantify the relative contribution of biological variability and measurement error across repeated assessments (Supplemental Material 1).

### Statistical Analysis

#### Agreement between quantitative gait measures and clinical assessment

To determine the association between hindlimb joint angles and TSCIS scores, we used the linear support vector machine (SVM) in the classification learner in MATLAB (The MathWorks, Inc.). The unit of analysis was gait cycles from one hind limb at one assessment session. Separate classification models were developed for the left and right hind limbs. Joint angle data from the pastern, ankle, knee, and hip throughout the gait cycle served as input features; corresponding TSCIS scores served as the classification labels (predictions). Before model training, joint angle data were standardized using z-score normalization based on the mean and standard deviation. Data were partitioned into training (80%) and testing (20%) sets. To prevent subject-level data leakage, all gait cycles from a given animal were assigned to the same partition (training or testing); the partition was stratified to balance treatment-group representation across folds.

The linear SVM was trained on the training dataset to identify optimal hyperplanes (decision boundaries) that best separate joint angle patterns associated with different TSCIS scores. Model performance was evaluated during model development via five-fold cross-validation within the training dataset. The final trained model was then applied to the testing dataset. Model performance was evaluated using classification accuracy and confusion matrices for both the training and testing datasets. For the testing dataset, macro-averaged precision, recall, F1 score, 95% confidence intervals (95% CI), and Cohen’s κ were also calculated. Following model training, the contribution of each joint to the TSCIS score classification was evaluated using the model coefficients (β weights). The absolute values of the coefficients were summed for each joint across the gait cycle to generate a single value of joint-level contribution.

#### Gait outcome comparisons

Gait outcomes were averaged for analysis. Linear mixed-effects models (Supplementary Material 2) were used to examine differences between treatment and control groups. This approach accounted for within-subject dependence arising from repeated measurements across limbs and time points. Fixed effects included treatment group, limb (treated vs. control), and time point (3–6 months), with appropriate interaction terms. Participant-specific random intercepts were included to account for between-subject variability. Model assumptions were assessed using Shapiro-Wilk tests and Q-Q plots of residuals; no violations were detected. Given the preliminary nature of this study and the constraints of the sheep model, no a priori sample size calculation was performed. Statistical significance was set at p<0.05. Linear mixed-effects models and discrete-outcome analyses were performed in SPSS (Version 29, IBM Corporation, Endicott, NY, USA).

#### Joint-space deviation

To identify between-group gait deviations by gait-cycle phase and by joint, we computed the Mahalanobis distance of each treated limb from the control reference distribution. At each point of the gait cycle, the pastern, ankle, knee, and hip angles were assembled into a four-dimensional feature vector. Mahalanobis distances were calculated using the average vector and covariance matrix estimated from the control group across the gait cycle. An analogous eight-dimensional Mahalanobis distance was calculated by combining joint angles from both limbs into a bilateral feature vector and comparing them with the corresponding pooled bilateral limbs in the control group. Distances were compared against 95th-percentile χ² reference values (χ²₄ = 3.08 for single-limb, four-joint distances; χ²₈ = 3.94 for the combined eight-joint distance) to identify the gait-cycle phases at which treated limbs deviated from the control group. To characterize the magnitude and temporal distribution of joint-specific deviations, the absolute standardized deviation (|z|) for each joint was calculated relative to the control group average and standard deviation and summarized across the gait cycle. Because the reference distribution was defined by a small control group, these analyses were considered exploratory.

## Results

Among 31 fetal lambs from 14 pregnant ewes, we created 28 models of spina bifida. The other three lambs served as normal controls. Twenty-five fetuses with the spina bifida defect underwent in-utero repair. Twenty-two treated lambs survived delivery; eight could not walk after birth and were excluded from gait analysis. The 14 ambulatory treated lambs and the three control lambs compose our study population (Figure 2). All treated lambs were grouped together for this analysis. One treated lamb was euthanized before 5-month evaluation because of disease progression.

**Figure 2.**
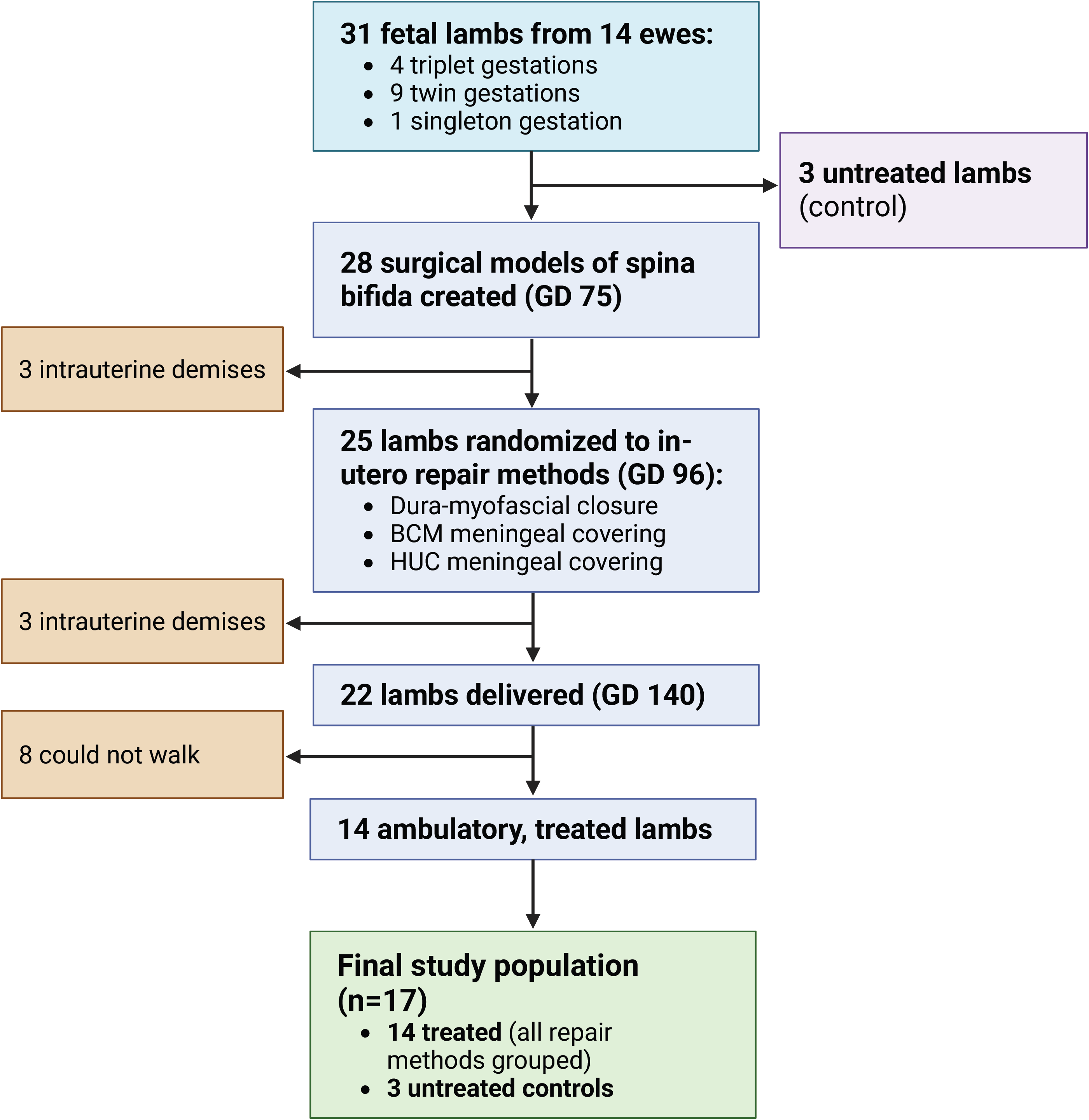
Flowchart of animals included in the study population. BCM, bovine collagen matrix graft; GD, gestational day; HUC, cryopreserved human umbilical cord graft.

### TSCIS assessment

Control lambs were consistently scored 6 in each hind limb at every time point; one had a right hindlimb score of 5 at 4 months. The 14 treated lambs had median [range] TSCIS scores of 5 [2–6] in both hind limbs at 3 and 4 months. These remained unchanged among the 13 surviving lambs at 5 and 6 months, except for right hindlimb scores of 5 [3–6] at 5 months. The lamb that was euthanized early had gait scores of 2 in both hind limbs at 3 and 4 months.

### Quantitative gait assessment and kinematic profiles

Using 3D motion capture data, we quantified spatiotemporal gait parameters across serial assessments (Table 1), and we calculated hindlimb sagittal hip, knee, ankle, and pastern joint angles across the gait cycle (Figure 3). Supplementary analysis confirmed the reliability of these repeated longitudinal measures: biological differences among lambs and their individual developmental trajectories were the predominant sources of variation; trial-related sources of measurement error contributed minimally to the observed measurements (Supplementary Material 1).

**Figure 3.**
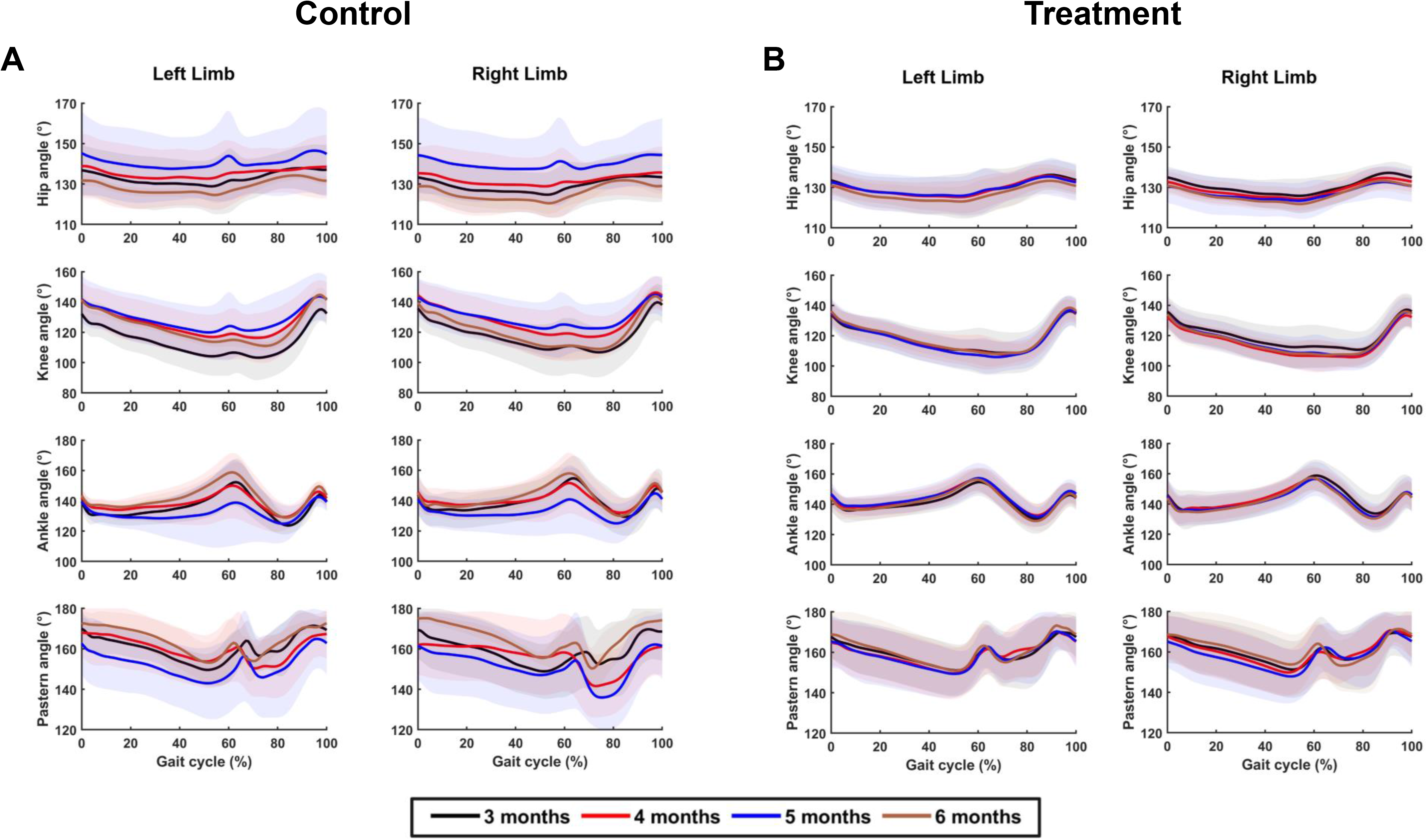
Hindlimb joint angles throughout the gait cycle across serial assessments. Descriptive sagittal plane hindlimb joint angles for (A) the control and (B) the treatment groups. Joint angles of the left and right limbs were plotted across the gait cycle. Panels are arranged from top to bottom as hip, knee, ankle, and pastern. Mean joint angles are shown as solid lines, with shaded regions representing standard deviations. Colors indicate assessment time points: black (3 months), red (4 months), blue (5 months), and brown (6 months). Extension was defined as 180°, with smaller angles indicating greater flexion.

Across groups, gait speed differed by time point in the front and hind limbs (both p=0.025). On pairwise comparison with Bonferroni correction, front- and hindlimb speeds were significantly slower at three months than at six months in both groups: front limb Δ=0.11 m/s, p=0.03 (Table 1A); hind limb Δ=0.11 m/s, p=0.03 (Table 1B). Stance and swing phases averaged 57.5 ± 2.9% and 42.5 ± 2.8% of the gait cycle, respectively, across time points and limbs. No other gait parameter differed by time point.

**Table 1A.**
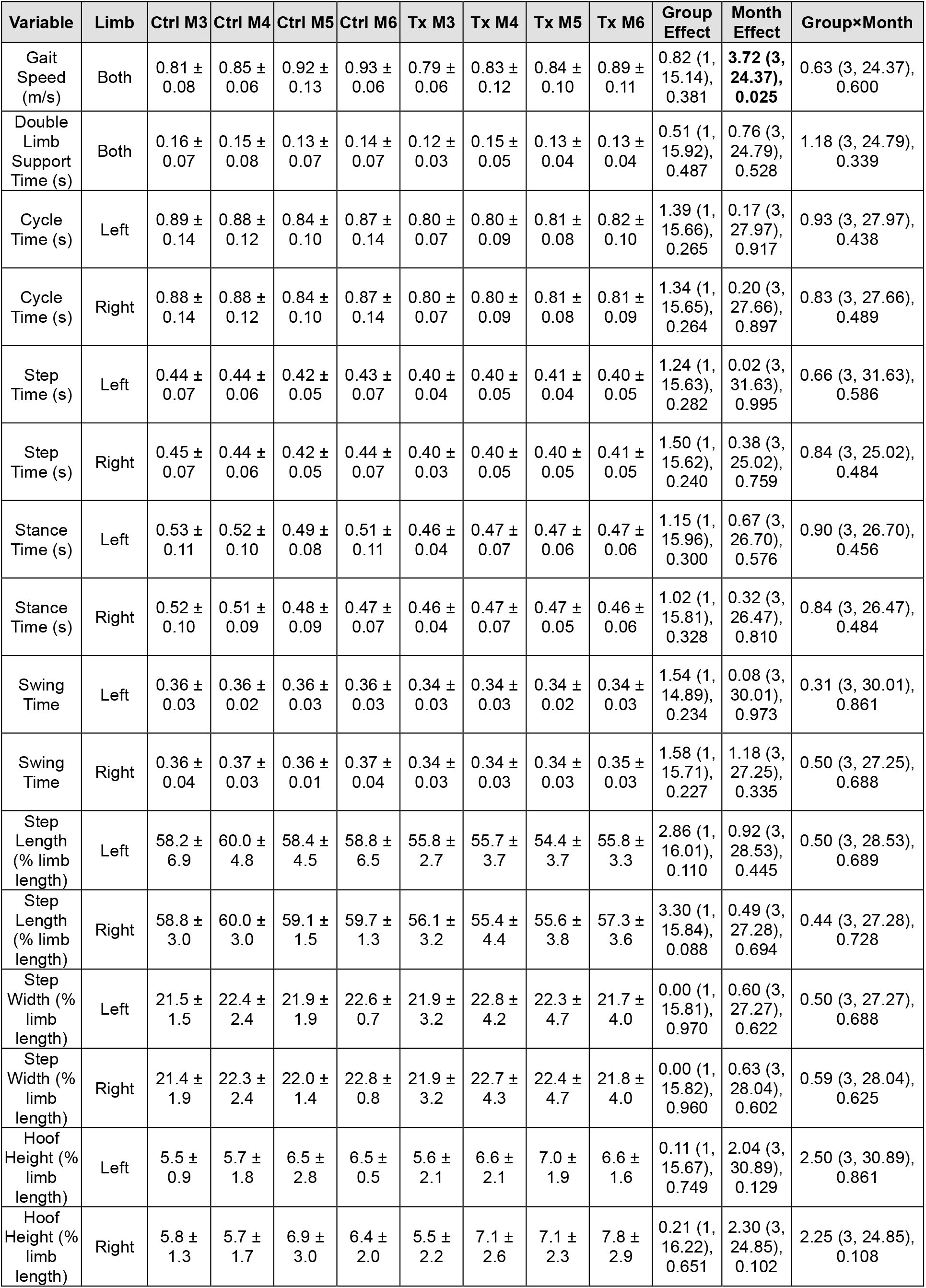
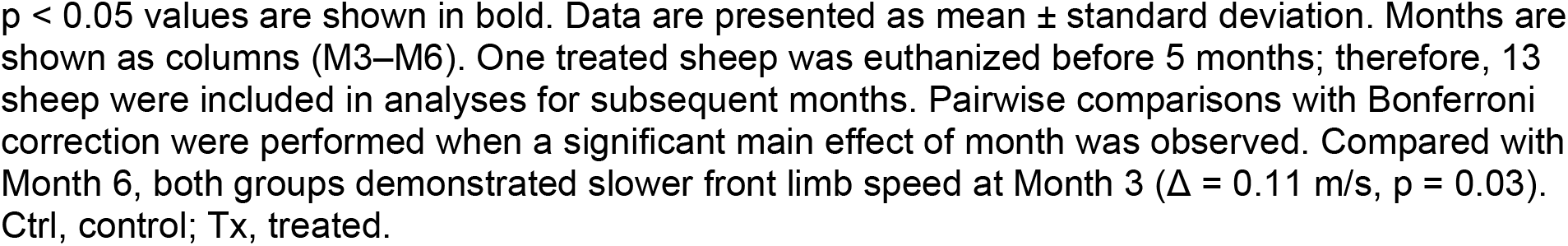
Descriptive and statistical analyses of gait outcomes for bilateral front limbs across all assessment times.

| Variable | Limb | Ctrl M3 | Ctrl M4 | Ctrl M5 | Ctrl M6 | Tx M3 | Tx M4 | Tx M5 | Tx M6 | Group Effect | Month Effect | Group×Month |
| --- | --- | --- | --- | --- | --- | --- | --- | --- | --- | --- | --- | --- |
| Gait Speed (m/s) | Both | 0.81 ± 0.08 | 0.85 ± 0.06 | 0.92 ± 0.13 | 0.93 ± 0.06 | 0.79 ± 0.06 | 0.83 ± 0.12 | 0.84 ± 0.10 | 0.89 ± 0.11 | 0.82 (1, 15.14), 0.381 | **3.72 (3, 24.37), 0.025** | 0.63 (3, 24.37), 0.600 |
| Double Limb Support Time (s) | Both | 0.16 ± 0.07 | 0.15 ± 0.08 | 0.13 ± 0.07 | 0.14 ± 0.07 | 0.12 ± 0.03 | 0.15 ± 0.05 | 0.13 ± 0.04 | 0.13 ± 0.04 | 0.51 (1, 15.92), 0.487 | 0.76 (3, 24.79), 0.528 | 1.18 (3, 24.79), 0.339 |
| Cycle Time (s) | Left | 0.89 ± 0.14 | 0.88 ± 0.12 | 0.84 ± 0.10 | 0.87 ± 0.14 | 0.80 ± 0.07 | 0.80 ± 0.09 | 0.81 ± 0.08 | 0.82 ± 0.10 | 1.39 (1, 15.66), 0.265 | 0.17 (3, 27.97), 0.917 | 0.93 (3, 27.97), 0.438 |
| Cycle Time (s) | Right | 0.88 ± 0.14 | 0.88 ± 0.12 | 0.84 ± 0.10 | 0.87 ± 0.14 | 0.80 ± 0.07 | 0.80 ± 0.09 | 0.81 ± 0.08 | 0.81 ± 0.09 | 1.34 (1, 15.65), 0.264 | 0.20 (3, 27.66), 0.897 | 0.83 (3, 27.66), 0.489 |
| Step Time (s) | Left | 0.44 ± 0.07 | 0.44 ± 0.06 | 0.42 ± 0.05 | 0.43 ± 0.07 | 0.40 ± 0.04 | 0.40 ± 0.05 | 0.41 ± 0.04 | 0.40 ± 0.05 | 1.24 (1, 15.63), 0.282 | 0.02 (3, 31.63), 0.995 | 0.66 (3, 31.63), 0.586 |
| Step Time (s) | Right | 0.45 ± 0.07 | 0.44 ± 0.06 | 0.42 ± 0.05 | 0.44 ± 0.07 | 0.40 ± 0.03 | 0.40 ± 0.05 | 0.40 ± 0.05 | 0.41 ± 0.05 | 1.50 (1, 15.62), 0.240 | 0.38 (3, 25.02), 0.759 | 0.84 (3, 25.02), 0.484 |
| Stance Time (s) | Left | 0.53 ± 0.11 | 0.52 ± 0.10 | 0.49 ± 0.08 | 0.51 ± 0.11 | 0.46 ± 0.04 | 0.47 ± 0.07 | 0.47 ± 0.06 | 0.47 ± 0.06 | 1.15 (1, 15.96), 0.300 | 0.67 (3, 26.70), 0.576 | 0.90 (3, 26.70), 0.456 |
| Stance Time (s) | Right | 0.52 ± 0.10 | 0.51 ± 0.09 | 0.48 ± 0.09 | 0.47 ± 0.07 | 0.46 ± 0.04 | 0.47 ± 0.07 | 0.47 ± 0.05 | 0.46 ± 0.06 | 1.02 (1, 15.81), 0.328 | 0.32 (3, 26.47), 0.810 | 0.84 (3, 26.47), 0.484 |
| Swing Time | Left | 0.36 ± 0.03 | 0.36 ± 0.02 | 0.36 ± 0.03 | 0.36 ± 0.03 | 0.34 ± 0.03 | 0.34 ± 0.03 | 0.34 ± 0.02 | 0.34 ± 0.03 | 1.54 (1, 14.89), 0.234 | 0.08 (3, 30.01), 0.973 | 0.31 (3, 30.01), 0.861 |
| Swing Time | Right | 0.36 ± 0.04 | 0.37 ± 0.03 | 0.36 ± 0.01 | 0.37 ± 0.04 | 0.34 ± 0.03 | 0.34 ± 0.03 | 0.34 ± 0.03 | 0.35 ± 0.03 | 1.58 (1, 15.71), 0.227 | 1.18 (3, 27.25), 0.335 | 0.50 (3, 27.25), 0.688 |
| Step Length (% limb length) | Left | 58.2 ± 6.9 | 60.0 ± 4.8 | 58.4 ± 4.5 | 58.8 ± 6.5 | 55.8 ± 2.7 | 55.7 ± 3.7 | 54.4 ± 3.7 | 55.8 ± 3.3 | 2.86 (1, 16.01), 0.110 | 0.92 (3, 28.53), 0.445 | 0.50 (3, 28.53), 0.689 |
| Step Length (% limb length) | Right | 58.8 ± 3.0 | 60.0 ± 3.0 | 59.1 ± 1.5 | 59.7 ± 1.3 | 56.1 ± 3.2 | 55.4 ± 4.4 | 55.6 ± 3.8 | 57.3 ± 3.6 | 3.30 (1, 15.84), 0.088 | 0.49 (3, 27.28), 0.694 | 0.44 (3, 27.28), 0.728 |
| Step Width (% limb length) | Left | 21.5 ± 1.5 | 22.4 ± 2.4 | 21.9 ± 1.9 | 22.6 ± 0.7 | 21.9 ± 3.2 | 22.8 ± 4.2 | 22.3 ± 4.7 | 21.7 ± 4.0 | 0.00 (1, 15.81), 0.970 | 0.60 (3, 27.27), 0.622 | 0.50 (3, 27.27), 0.688 |
| Step Width (% limb length) | Right | 21.4 ± 1.9 | 22.3 ± 2.4 | 22.0 ± 1.4 | 22.8 ± 0.8 | 21.9 ± 3.2 | 22.7 ± 4.3 | 22.4 ± 4.7 | 21.8 ± 4.0 | 0.00 (1, 15.82), 0.960 | 0.63 (3, 28.04), 0.602 | 0.59 (3, 28.04), 0.625 |
| Hoof Height (% limb length) | Left | 5.5 ± 0.9 | 5.7 ± 1.8 | 6.5 ± 2.8 | 6.5 ± 0.5 | 5.6 ± 2.1 | 6.6 ± 2.1 | 7.0 ± 1.9 | 6.6 ± 1.6 | 0.11 (1, 15.67), 0.749 | 2.04 (3, 30.89), 0.129 | 2.50 (3, 30.89), 0.861 |
| Hoof Height (% limb length) | Right | 5.8 ± 1.3 | 5.7 ± 1.7 | 6.9 ± 3.0 | 6.4 ± 2.0 | 5.5 ± 2.2 | 7.1 ± 2.6 | 7.1 ± 2.3 | 7.8 ± 2.9 | 0.21 (1, 16.22), 0.651 | 2.30 (3, 24.85), 0.102 | 2.25 (3, 24.85), 0.108 |

p < 0.05 values are shown in bold. Data are presented as mean ± standard deviation. Months are shown as columns (M3–M6). One treated sheep was euthanized before 5 months; therefore, 13 sheep were included in analyses for subsequent months. Pairwise comparisons with Bonferroni correction were performed when a significant main effect of month was observed. Compared with Month 6, both groups demonstrated slower front limb speed at Month 3 ( $\Delta = 0.11$ m/s, $p = 0.03$ ). Ctrl, control; Tx, treated.

**Table 1B.**
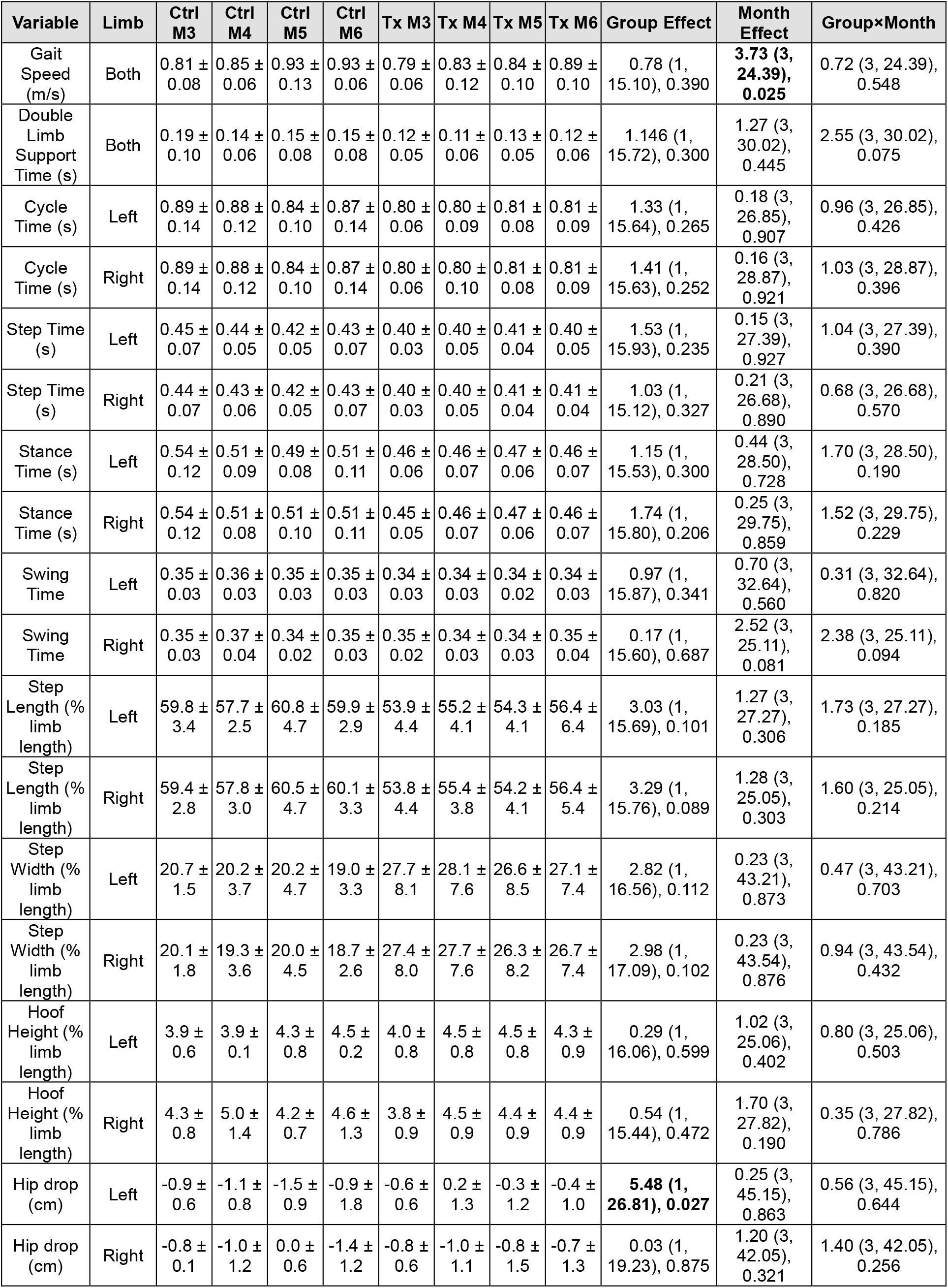
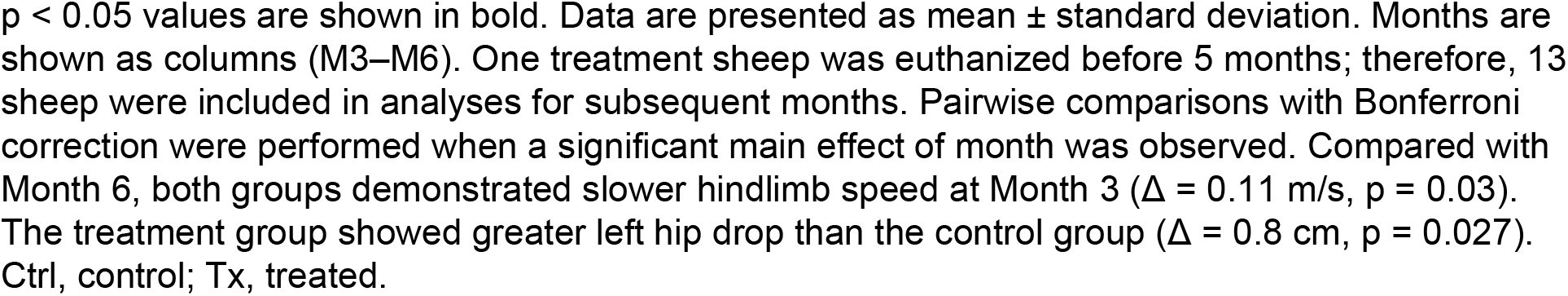
Descriptive and statistical analyses of gait outcomes for bilateral hind limbs across all assessment times.

| Variable | Limb | Ctrl M3 | Ctrl M4 | Ctrl M5 | Ctrl M6 | Tx M3 | Tx M4 | Tx M5 | Tx M6 | Group Effect | Month Effect | Group×Month |
| --- | --- | --- | --- | --- | --- | --- | --- | --- | --- | --- | --- | --- |
| Gait Speed (m/s) | Both | 0.81 ± 0.08 | 0.85 ± 0.06 | 0.93 ± 0.13 | 0.93 ± 0.06 | 0.79 ± 0.06 | 0.83 ± 0.12 | 0.84 ± 0.10 | 0.89 ± 0.10 | 0.78 (1, 15.10), 0.390 | **3.73 (3, 24.39), 0.025** | 0.72 (3, 24.39), 0.548 |
| Double Limb Support Time (s) | Both | 0.19 ± 0.10 | 0.14 ± 0.06 | 0.15 ± 0.08 | 0.15 ± 0.08 | 0.12 ± 0.05 | 0.11 ± 0.06 | 0.13 ± 0.05 | 0.12 ± 0.06 | 1.146 (1, 15.72), 0.300 | 1.27 (3, 30.02), 0.445 | 2.55 (3, 30.02), 0.075 |
| Cycle Time (s) | Left | 0.89 ± 0.14 | 0.88 ± 0.12 | 0.84 ± 0.10 | 0.87 ± 0.14 | 0.80 ± 0.06 | 0.80 ± 0.09 | 0.81 ± 0.08 | 0.81 ± 0.09 | 1.33 (1, 15.64), 0.265 | 0.18 (3, 26.85), 0.907 | 0.96 (3, 26.85), 0.426 |
| Cycle Time (s) | Right | 0.89 ± 0.14 | 0.88 ± 0.12 | 0.84 ± 0.10 | 0.87 ± 0.14 | 0.80 ± 0.06 | 0.80 ± 0.10 | 0.81 ± 0.08 | 0.81 ± 0.09 | 1.41 (1, 15.63), 0.252 | 0.16 (3, 28.87), 0.921 | 1.03 (3, 28.87), 0.396 |
| Step Time (s) | Left | 0.45 ± 0.07 | 0.44 ± 0.05 | 0.42 ± 0.05 | 0.43 ± 0.07 | 0.40 ± 0.03 | 0.40 ± 0.05 | 0.41 ± 0.04 | 0.40 ± 0.05 | 1.53 (1, 15.93), 0.235 | 0.15 (3, 27.39), 0.927 | 1.04 (3, 27.39), 0.390 |
| Step Time (s) | Right | 0.44 ± 0.07 | 0.43 ± 0.06 | 0.42 ± 0.05 | 0.43 ± 0.07 | 0.40 ± 0.03 | 0.40 ± 0.05 | 0.41 ± 0.04 | 0.41 ± 0.04 | 1.03 (1, 15.12), 0.327 | 0.21 (3, 26.68), 0.890 | 0.68 (3, 26.68), 0.570 |
| Stance Time (s) | Left | 0.54 ± 0.12 | 0.51 ± 0.09 | 0.49 ± 0.08 | 0.51 ± 0.11 | 0.46 ± 0.06 | 0.46 ± 0.07 | 0.47 ± 0.06 | 0.46 ± 0.07 | 1.15 (1, 15.53), 0.300 | 0.44 (3, 28.50), 0.728 | 1.70 (3, 28.50), 0.190 |
| Stance Time (s) | Right | 0.54 ± 0.12 | 0.51 ± 0.08 | 0.51 ± 0.10 | 0.51 ± 0.11 | 0.45 ± 0.05 | 0.46 ± 0.07 | 0.47 ± 0.06 | 0.46 ± 0.07 | 1.74 (1, 15.80), 0.206 | 0.25 (3, 29.75), 0.859 | 1.52 (3, 29.75), 0.229 |
| Swing Time | Left | 0.35 ± 0.03 | 0.36 ± 0.03 | 0.35 ± 0.03 | 0.35 ± 0.03 | 0.34 ± 0.03 | 0.34 ± 0.03 | 0.34 ± 0.02 | 0.34 ± 0.03 | 0.97 (1, 15.87), 0.341 | 0.70 (3, 32.64), 0.560 | 0.31 (3, 32.64), 0.820 |
| Swing Time | Right | 0.35 ± 0.03 | 0.37 ± 0.04 | 0.34 ± 0.02 | 0.35 ± 0.03 | 0.35 ± 0.02 | 0.34 ± 0.03 | 0.34 ± 0.03 | 0.35 ± 0.04 | 0.17 (1, 15.60), 0.687 | 2.52 (3, 25.11), 0.081 | 2.38 (3, 25.11), 0.094 |
| Step Length (% limb length) | Left | 59.8 ± 3.4 | 57.7 ± 2.5 | 60.8 ± 4.7 | 59.9 ± 2.9 | 53.9 ± 4.4 | 55.2 ± 4.1 | 54.3 ± 4.1 | 56.4 ± 6.4 | 3.03 (1, 15.69), 0.101 | 1.27 (3, 27.27), 0.306 | 1.73 (3, 27.27), 0.185 |
| Step Length (% limb length) | Right | 59.4 ± 2.8 | 57.8 ± 3.0 | 60.5 ± 4.7 | 60.1 ± 3.3 | 53.8 ± 4.4 | 55.4 ± 3.8 | 54.2 ± 4.1 | 56.4 ± 5.4 | 3.29 (1, 15.76), 0.089 | 1.28 (3, 25.05), 0.303 | 1.60 (3, 25.05), 0.214 |
| Step Width (% limb length) | Left | 20.7 ± 1.5 | 20.2 ± 3.7 | 20.2 ± 4.7 | 19.0 ± 3.3 | 27.7 ± 8.1 | 28.1 ± 7.6 | 26.6 ± 8.5 | 27.1 ± 7.4 | 2.82 (1, 16.56), 0.112 | 0.23 (3, 43.21), 0.873 | 0.47 (3, 43.21), 0.703 |
| Step Width (% limb length) | Right | 20.1 ± 1.8 | 19.3 ± 3.6 | 20.0 ± 4.5 | 18.7 ± 2.6 | 27.4 ± 8.0 | 27.7 ± 7.6 | 26.3 ± 8.2 | 26.7 ± 7.4 | 2.98 (1, 17.09), 0.102 | 0.23 (3, 43.54), 0.876 | 0.94 (3, 43.54), 0.432 |
| Hoof Height (% limb length) | Left | 3.9 ± 0.6 | 3.9 ± 0.1 | 4.3 ± 0.8 | 4.5 ± 0.2 | 4.0 ± 0.8 | 4.5 ± 0.8 | 4.5 ± 0.8 | 4.3 ± 0.9 | 0.29 (1, 16.06), 0.599 | 1.02 (3, 25.06), 0.402 | 0.80 (3, 25.06), 0.503 |
| Hoof Height (% limb length) | Right | 4.3 ± 0.8 | 5.0 ± 1.4 | 4.2 ± 0.7 | 4.6 ± 1.3 | 3.8 ± 0.9 | 4.5 ± 0.9 | 4.4 ± 0.9 | 4.4 ± 0.9 | 0.54 (1, 15.44), 0.472 | 1.70 (3, 27.82), 0.190 | 0.35 (3, 27.82), 0.786 |
| Hip drop (cm) | Left | -0.9 ± 0.6 | -1.1 ± 0.8 | -1.5 ± 0.9 | -0.9 ± 1.8 | -0.6 ± 0.6 | 0.2 ± 1.3 | -0.3 ± 1.2 | -0.4 ± 1.0 | **5.48 (1, 26.81), 0.027** | 0.25 (3, 45.15), 0.863 | 0.56 (3, 45.15), 0.644 |
| Hip drop (cm) | Right | -0.8 ± 0.1 | -1.0 ± 1.2 | 0.0 ± 0.6 | -1.4 ± 1.2 | -0.8 ± 0.6 | -1.0 ± 1.1 | -0.8 ± 1.5 | -0.7 ± 1.3 | 0.03 (1, 19.23), 0.875 | 1.20 (3, 42.05), 0.321 | 1.40 (3, 42.05), 0.256 |

p < 0.05 values are shown in bold. Data are presented as mean ± standard deviation. Months are shown as columns (M3–M6). One treatment sheep was euthanized before 5 months; therefore, 13 sheep were included in analyses for subsequent months. Pairwise comparisons with Bonferroni correction were performed when a significant main effect of month was observed. Compared with Month 6, both groups demonstrated slower hindlimb speed at Month 3 ( $\Delta = 0.11$ m/s, $p = 0.03$ ). The treatment group showed greater left hip drop than the control group ( $\Delta = 0.8$ cm, $p = 0.027$ ). Ctrl, control; Tx, treated.

### Agreement between TSCIS scores and joint angle profile

Across 17 animals, 1,632 left and 1,661 right hindlimb gait cycles were separated into training and testing datasets: 1,306 training and 326 testing samples for the left limb, 1,329 training and 332 testing samples for the right limb. TSCIS scores of 5 or 6 accounted for 76.5% and 75% of the left and right hindlimb evaluations, respectively. Lower-function scores (2-3) represented 6.2% and 16.2% of left and right evaluations, respectively (Figure 4).

**Figure 4.**
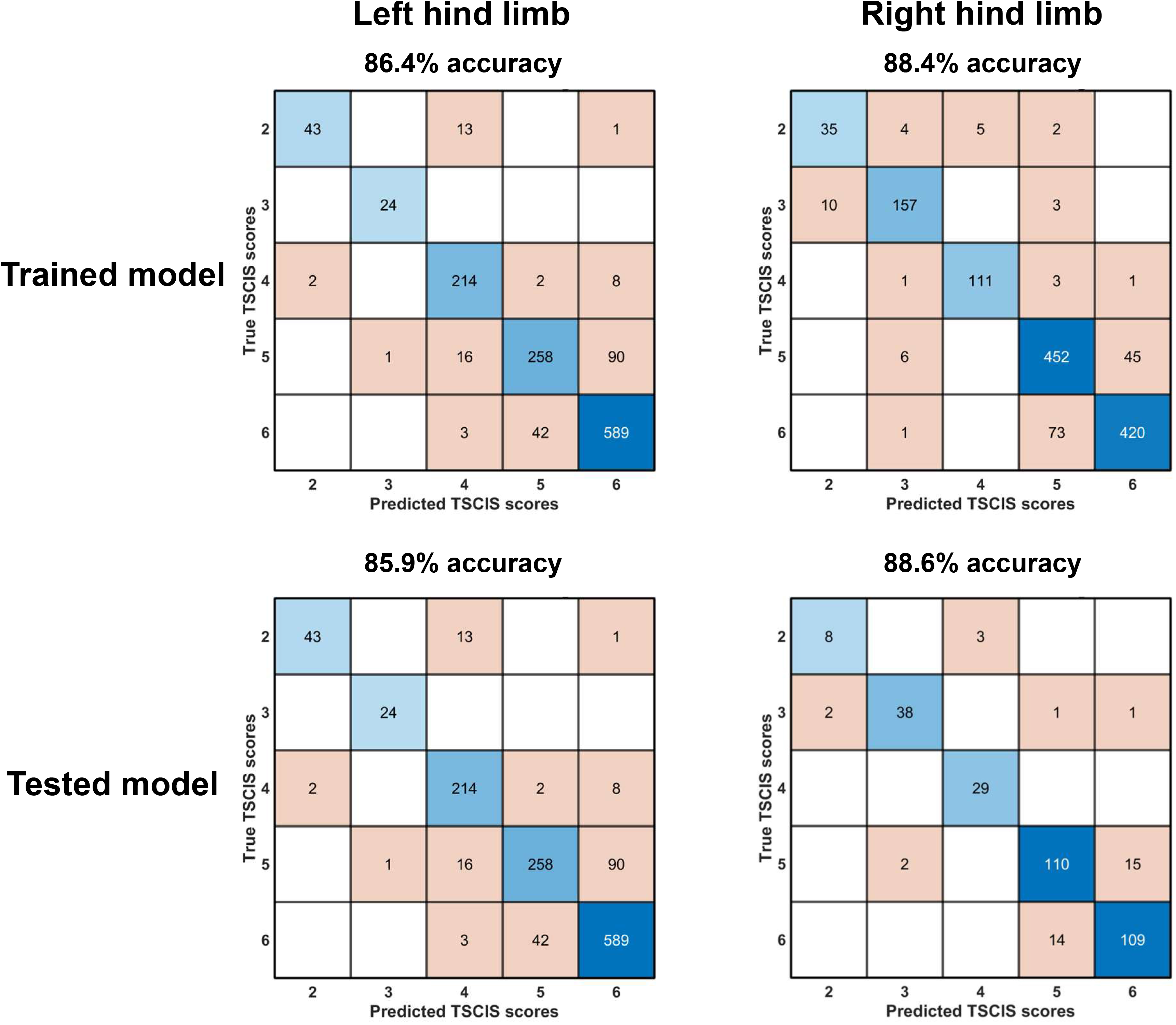
Quantitative limb joint angle profiles show strong agreement with clinical assessments. Confusion matrices show agreement between left and right hindlimb joint angle profiles and clinician-assessed Texas Spinal Cord Injury Scale (TSCIS) scores. Top matrices show model performance on the training dataset. Bottom matrices show model performance on the held-out testing dataset. Each row represents the clinician-rated TSCIS scores (True TSCIS scores), and each column represents the predicted TSCIS scores based on hindlimb sagittal plane joint angle profiles. Blue shading indicates correct classification along the diagonal, representing agreement between predicted and true TSCIS scores. Orange shading indicates misclassifications (disagreement between predicted and true TSCIS scores). Darker color intensity reflects stronger agreement/disagreement.

Per-class confusion matrices are shown in Figure 4. Five-fold cross-validated training accuracy was 86.4% for the left and 88.4% for the right limb. Held-out test accuracy was 85.9% for the left and 88.6% for the right limb. Macro-averaged precision, recall, and F1 across TSCIS classes were 89.4%, 88.7%, and 88.8% for the left limb model and 88.2%, 87.7%, and 87.8% for the right limb model, respectively. The Cohen’s κ was 0.78 (95% CI=0.73–0.84) and 0.84 (95% CI=0.79–0.88) for the left and right models, respectively.

Misclassifications occurred most often between adjacent high-functioning categories: 28.3% of TSCIS 5 samples were classified as TSCIS 4 or 6, and 9.5% of TSCIS 6 samples were classified as TSCIS 4 or 5. Samples with TSCIS ≤ 4 were classified with 88.5% accuracy. The pastern joint angle contributed most to the TSCIS classification in both the left and right limb models, accounting for 31.3% and 35.7%, respectively, of the contribution, followed by ankle, knee, and hip joints (Figure 5).

**Figure 5.**
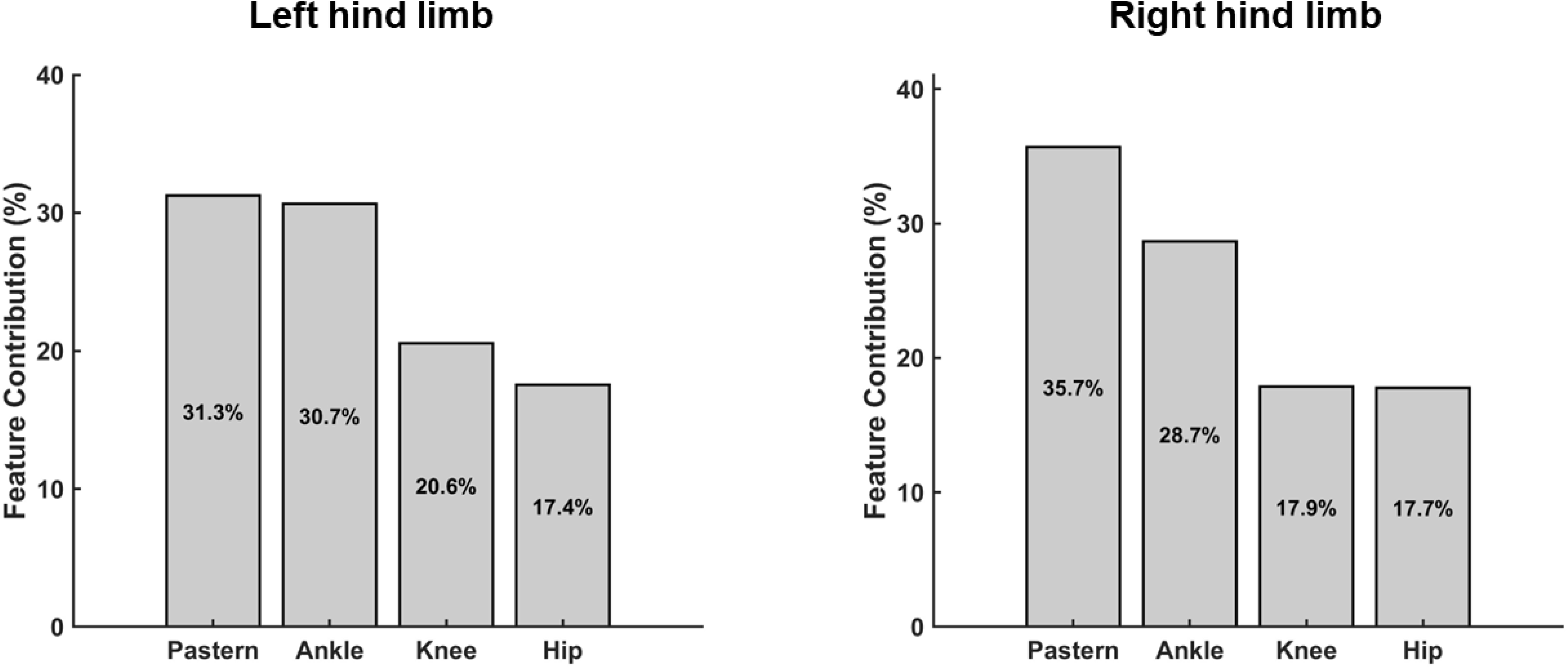
Joint feature contribution to the Texas Spinal Cord Injury Scale (TSCIS) score classification. Bar graphs depict the feature contribution of each hindlimb joint angle to the TSCIS score classification for the left and right hind limbs. Percentage values within each bar represent the relative contribution of each joint to the classification.

### Gait differences between groups

On exploratory analysis, left hindlimb hip drop differed by group: treated lambs exhibited a greater left hip drop than control lambs (−0.3 ± 1.1 cm vs. −1.1 ± 0.8 cm, Δ=0.8 cm, p=0.027) (Table 1B). No other between-group differences were statistically significant.

### Localization of gait deviations

In the exploratory joint-space deviation analysis, treated lambs’ gait deviated most from control lambs’ during the initial swing phase (left limb) and the mid-stance phase (right limb). The mean Mahalanobis distance peaked at approximately 71% of the gait cycle for the left limb (2.04), 24% for the right limb (1.99), and 71% for the combined bilateral analysis (3.94); however, these values were within the 95% confidence interval of the reference distribution (Figure 6A). At the joint level, the left limb demonstrated the largest average standardized deviations at the knee (mean |z| = 1.26) and hip (1.15); the right limb showed the largest deviations at the ankle (1.10) and hip (1.22) (Figure 6B). Left limb deviations were concentrated in the pastern during mid-swing phase, while the right limb exhibited ankle deviations during the early stance phase and increased pastern deviations during mid-swing phase (Figure 6C). Gait deviation results for each treated lamb are presented in Supplementary Material 3.

**Figure 6.**
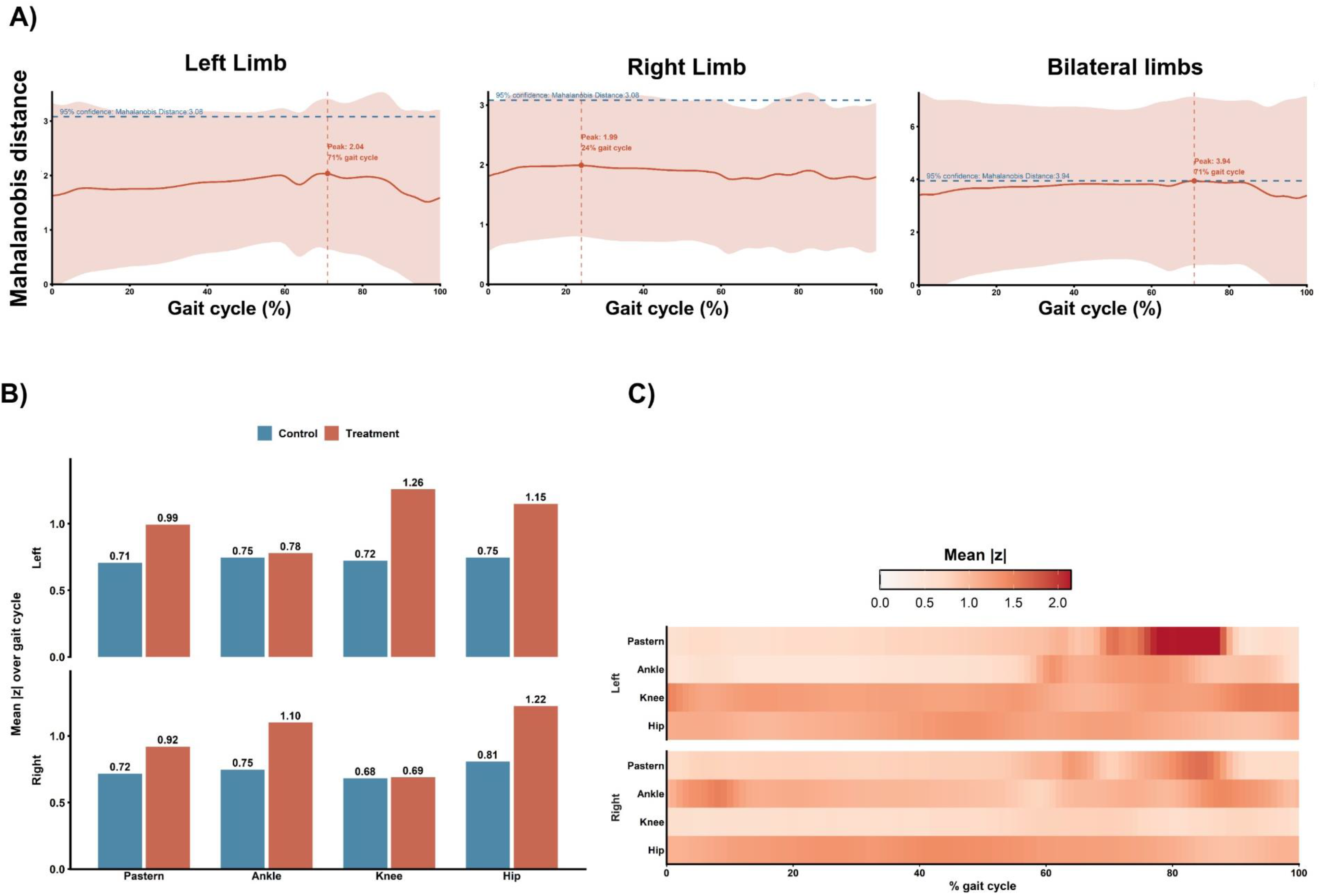
Mahalanobis distance and standardized joint angle profiles. (A) Left, right, and bilateral limb Mahalanobis distances across the gait cycle. Solid lines represent the mean treatment group Mahalanobis distance relative to the control group. Dashed horizontal lines indicate the 95% χ² reference threshold, and vertical lines indicate the gait cycle location of the peak treatment group Mahalanobis distance. (B) Bar graphs show the mean absolute standardized deviation (|z|) for each joint in the Control (blue) and Treatment (red) groups across all eight joints (four per limb). Joints with larger mean |z| values contribute more strongly to the overall Mahalanobis distance. (C) Heat map shows the treatment group mean absolute standardized deviation (|z|) across the gait cycle for each joint and limb.

## Discussion

We implemented a 3D marker-based motion capture system to objectively quantify spatiotemporal gait parameters and hindlimb joint angles over serial assessments in an established sheep model of in-utero spina bifida repair, and we confirmed the reliability of the repeated measurements. Using this system, we detected changes in gait parameters over time, and exploratory analyses quantified differences in gait symmetry between treated lambs and controls. Hindlimb kinematic profiles demonstrated accuracy above the majority-class baseline in classifying clinician-assigned TSCIS scores, which supports the functional relevance of these gait measures.

To our knowledge, this study is the first to apply marker-based 3D motion capture to the sheep model of in-utero spina bifida repair and to serially track quantitative spatiotemporal and multi-joint kinematic parameters to 6 months of age. The sheep model has long been essential to the development and testing of in-utero repair techniques.^19–23^ However, prior studies in this model characterized postnatal motor function almost exclusively through ordinal locomotor rating scales.^15–17,23–26^ These scales compress complex locomotor behavior into a few discrete categories and remain inherently examiner-dependent, which limits sensitivity for evaluating or comparing treatments. In contrast, quantitative biomechanical gait analysis enables continuous, reproducible, and high-dimensional measures of locomotor function, providing substantially greater resolution for characterizing locomotor behavior and detecting subtle abnormalities that may not be apparent during visual assessment. In leveraging these capabilities, our study seeks to extend prior work in the sheep model toward continuous, reproducible, and developmentally resolved measures of ambulatory recovery after in-utero repair.

An important strength of marker-based motion capture is the breadth and dimensionality of information generated during walking. Movement is quantified continuously across time while simultaneously capturing the dynamic interactions among multiple joints, limbs, and phases of gait. This multidimensional biomechanical dataset enables the evaluation of locomotor patterns at a level of temporal and spatial precision beyond the limits of human observation, even for experienced clinicians. Consequently, quantitative gait analysis can identify compensatory movement strategies and evolving abnormalities before they may be observable in the clinic. These capabilities are particularly relevant in spina bifida, where impairment often manifests as complex, evolving compensatory gait strategies rather than isolated deficits at a single joint or anatomical level.^35–38^ In the present study, quantitative analysis detected a between-group difference in left hindlimb hip drop that categorical scoring did not capture. Joint-space deviation analysis identified the points of greatest separation from the control reference at the initial swing and mid-stance phases in the left and right limbs, respectively. The joints driving the deviation differed by side, potentially indicating a clinically meaningful gait asymmetry. However, these exploratory findings rest on only three control animals and should be regarded as hypothesis-generating until confirmed in a larger cohort.

By using supervised machine learning to classify gait, our study is connected to the emerging research applying machine learning to gait analysis.^39^ Support vector machines and related classifiers have been applied in Parkinson’s disease, cerebral palsy, autism spectrum disorder, stroke and traumatic brain injury, with reported accuracies ranging from 80-95% depending on the condition, feature set, and validation scheme.^40–44^ Notably, machine-learning models trained on kinematic features have reproduced examiner-assigned ordinal gait ratings with agreement comparable to that between human raters.^44^ Our per-cycle classification accuracies of 85.9% and 88.6% fall within this reported range despite using only sagittal-plane joint-angle waveforms as input features. Nonetheless, these accuracies should be interpreted relative to the majority-class baseline and the class distribution of our sample. To our knowledge, however, ours is the first study to apply such a classification framework to a large-animal fetal-surgery model, linking objectively measured hindlimb kinematics to a clinically used ambulatory scale.

Our study’s main strength is its use of marker-based motion capture—considered the “gold standard” method for biomechanical gait measurement^28,45^—in a well-established animal model of in-utero spina bifida repair. Animal surgeries and TSCIS assessments were conducted under a prospective research protocol with consistent data collection. Also, assessing gait on a mat walkway yielded more ecologically valid results than assessments on a treadmill. Nonetheless, the study was limited by its sample size, which hampered comparisons between groups and among surgical techniques. The timing of assessments was driven by factors specific to the animal model—such as housing and caring for them in a research facility—rather than by clinically relevant time points. Finally, treated lambs that could not walk after delivery were excluded.

Future studies in larger, powered cohorts are needed to confirm exploratory findings and to allow comparisons among repair techniques. A larger cohort would also permit leave-one-subject-out cross-validation as the primary evaluation of the classification models, reducing the influence of the pseudoreplication inherent in analyzing many gait cycles from a small number of animals. Expanding the kinematic model to include frontal- and transverse-plane motion, together with kinetic measures from instrumented force platforms, would more fully characterize the compensatory and asymmetric strategies that categorical scoring cannot resolve. Finally, relating these quantitative gait signatures to spinal cord histology, neuro-segmental level, and longer-term functional outcomes would clarify the biological basis of the model’s classifications and strengthen their validity as surrogates for ambulatory recovery.

The substantial proportion of children who remain unable to ambulate independently even after in-utero repair underscores the need for sensitive, objective, and reproducible measures of ambulatory function to evaluate emerging repair strategies.^1^ The framework demonstrated here—quantitative motion capture combined with machine-learning classification against a clinically relevant scale—provides such a measure in a controlled preclinical setting. Emerging markerless and portable motion-capture approaches^46^ may lower the barrier to acquiring high-dimensional gait data and, in time, support translation of this approach to the clinic.

## Supporting information

Supplement 1 - Reliability Analysis

Supplement 2 - LMM Equation

Supplement 3 - Mahalanobis Distance

## Data availability

Raw motion-capture data (C3D files), processed joint-angle waveforms and spatiotemporal parameters (CSV), TSCIS scores, and trained SVM model objects will be deposited in a public repository (Zenodo) with a citable DOI upon acceptance. Analysis code, including the Visual3D processing pipeline (.cmo and .v3s files), MATLAB SVM training and evaluation scripts, and the linear mixed-effects model syntax, will be released on GitHub with a commit-pinned versioned release linked from the data record. Animal-level metadata required to reproduce subject-level partitioning is included with the deposit. Any data not deposited due to file-size or licensing constraints are available from the corresponding author upon reasonable request.

## Author contribution statement

All listed authors contributed substantially to the study design or to the acquisition, analysis, or interpretation of data, as follows. **TYC:** data analysis and interpretation; manuscript drafting and review. **SP:** data collection and analysis; manuscript review. **PZ:** data analysis and interpretation, manuscript drafting and review. **LFF:** data collection; manuscript review. **DL:** data analysis and interpretation; manuscript review. **RP:** study design and oversight; animal surgeries; data analysis and interpretation; manuscript drafting and review. **LKM:** study design and oversight; data collection, analysis and interpretation; manuscript review.

## Ethical approval statement

This study was conducted under the following protocols approved by the UTHealth Houston Institutional Animal Care and Use Committee: protocol AWC-20-0149, approved May 11, 2021, and protocol AWC-23-0115, approved January 10, 2024. Animal care was conducted in compliance with the Guide for the Care and Use of Laboratory Animals.

## Patient consent statement

Patient consent was not applicable for this preclinical study in an animal model.

## Acknowledgements

Jonathan S. Feinberg, PhD, provided editorial support in preparing the manuscript.

