## Supplement 1 - Reliability Analysis for "Quantitative Gait Analysis in a Sheep Model of In-Utero Spina Bifida Repair"

**Supplementary Material 1. Reliability analysis.**

We analyzed the reliability of repeated longitudinal measures by applying Generalizability Theory (G-theory), a comprehensive framework for decomposing measurement variance into its constituent sources and evaluating how modifications to the measurement protocol influence reliability.[1]

Using Generalizability Theory analysis, we (1) quantified the relative contributions of lamb, developmental assessment month, repeated gait trials, and their interactions to the variability in spatiotemporal gait parameters, and (2) determined the minimum number of gait cycles required to obtain dependable gait measurements for future longitudinal studies in ovine models of spina bifida.

### A METHODS

#### A.1 Subjects and Data Collection Protocol

Longitudinal gait data were collected from three control lambs assessed monthly from months 3-6 after birth. At each assessment, animals traversed a gait analysis walkway while 10 successful gait cycles were collected under consistent experimental conditions. In total, 38 gait outcome measures were evaluated for each gait cycle, including bilateral limb-specific variables and whole-animal measures.

Each gait cycle was treated as an independent repeated observation within an assessment session. Consequently, the dataset consisted of repeated measurements nested within lambs across multiple longitudinal assessment months, providing the hierarchical structure required for Generalizability Theory analysis.

#### A.2 Generalizability Theory Analysis (G-Study)

The reliability of each gait outcome was evaluated using Generalizability Theory (G-theory) to quantify the relative contribution of biological variability and measurement error across repeated gait assessments. A fully crossed random-effects design was specified with Lamb, Month, and Trial as facets of measurement:

$$Y_{lmt} = \mu + L + M + T + LM + LT + MT + \varepsilon, \quad \text{Equation 1}$$

where L represents the effect of lamb, M the assessment month, T the gait trial, LT, MT, and LM are their two-way interactions, and $\epsilon$ the residual variance, which also incorporates the unmodeled three-way interaction (Lamb × Month × Trial).

Accordingly, the total variance was partitioned as

$$\sigma_Y^2 = \sigma_L^2 + \sigma_M^2 + \sigma_T^2 + \sigma_{LM}^2 + \sigma_{LT}^2 + \sigma_{MT}^2 + \sigma_E^2, \quad \text{Equation 2}$$

where $\sigma_L^2$ represents biological differences among lambs, $\sigma_M^2$ reflects variability attributable to developmental assessment month, $\sigma_T^2$ represents trial-to-trial variability, $\sigma_{LM}^2$ quantifies differences in developmental trajectories among lambs, $\sigma_{LT}^2$ reflects differential trial-to-trial variability among lambs, $\sigma_{MT}^2$ captures changes in trial effects across assessment months, and $\sigma_E^2$ represents residual variance, including the unmodeled three-way interaction and measurement error.

Variance components were estimated separately for each gait outcome using restricted maximum likelihood implemented in the gtheory package in R. The proportion of total variance attributable to each source was calculated to characterize the relative contribution of biological variability, longitudinal development, repeated gait trials, and residual measurement error.

#### A.3 Decision Study Analysis (D-Study)

A D-study was subsequently performed to determine the minimum number of gait trials required to achieve acceptable measurement reliability in future longitudinal studies.

The assessment months were treated as a fixed facet because they represent a predefined longitudinal protocol rather than a random sample of developmental occasions. Accordingly, the universe score was defined as the expected gait measurement for each lamb under this fixed assessment schedule. Variance attributable to Lamb and the Lamb × Month interaction was therefore interpreted as genuine biological variation, reflecting individual differences in gait and developmental trajectories, whereas Trial and Lamb × Trial represented measurement variability arising from repeated gait cycles and were incorporated into the D-study error variance.

Therefore, the dependability coefficient (Φ), representing absolute agreement, was calculated as

$$\Phi(n_T) = \frac{\sigma_L^2}{\sigma_L^2 + \frac{\sigma_T^2 + \sigma_{LT}^2 + \sigma_E^2}{n_T}}, \qquad \text{Equation 3}$$

where $\sigma_L^2$ is the variance attributable to lamb, $\sigma_T^2$ is the trial variance, $\sigma_{LT}^2$ is the Lamb × Trial interaction variance, $\sigma_{MLT,\epsilon}^2$ is the residual variance, $n_T$ is the number of gait trials averaged. Dependability coefficients were computed for $n_T$ from 1 through 10 gait trials for every gait outcome. The minimum number of trials required to achieve predetermined reliability thresholds was determined for each outcome measure.

A target dependability coefficient of Φ ≥ 0.80 was considered adequate for repeated biomechanical assessments in the present study, in keeping with conventional reliability benchmarks in both psychometric and biomedical research, whereby Φ values of approximately 0.70 are considered acceptable for exploratory or group-level investigations, 0.80 or greater indicate good dependability for most clinical and biomechanical research applications, and 0.90 or greater reflect excellent reliability for high-stakes individual-level decisions.[2,3] Accordingly, and the minimum number of trials required to achieve this threshold was determined using D-study analyses.

As such, for each outcome measure, the minimum number of gait cycles required was determined by identifying the smallest number of averaged trials that achieved a dependability coefficient ($\Phi$) of at least 0.80, which was considered indicative of acceptable reliability.

#### A.4 Statistical Rationale

Generalizability Theory partitions measurement variability into biologically meaningful and measurement-related sources. In the present study, the objective was to determine the number of gait cycles required to obtain stable gait estimates during a fixed longitudinal developmental protocol, rather than to optimize the timing of developmental assessments. Consequently, developmental differences among lambs across assessment months were interpreted as genuine biological variation, whereas variability attributable to repeated gait trials was treated as measurement uncertainty that could be reduced by averaging multiple gait cycles. Accordingly, only trial-related variance components contributed to the D-study error term.

### B RESULTS

#### B.1 Generalizability Study

The G-study partitioned the variance of each gait outcome into components attributable to Lamb, Month, Trial, their two-way interactions, and residual error (Table S1).

Across most gait outcomes, between-lamb variability accounted for the largest proportion of the total variance, particularly for temporal and spatial gait parameters. For temporal variables, the Lamb component consistently explained approximately 40.6–60.5% of the total variance, with cycle duration, stance time, step time, and double-limb support time exhibiting the greatest between-lamb variability (51.3–60.5%). Similarly, stride length and step length demonstrated substantial Lamb variance (44.3–58.6%), whereas hind limb step width exhibited the greatest Lamb contribution among the step width measures (63.4–66.6%).

The Lamb × Month interaction represented the second largest source of variability for most gait parameters, accounting for approximately 7.3–27.7% of the total variance in temporal and spatial outcomes. This interaction was particularly prominent for pelvic kinematics, explaining 60.1% and 63.4% of the variance in right and left hip drop, respectively, and for right hind hoof height, where it accounted for 50.4% of the total variance. These findings indicate that developmental changes differed among individual lambs throughout the longitudinal study.

The contribution of Month as a main effect was generally negligible for temporal gait variables, typically accounting for 0–1.2% of the variance. In contrast, Month explained a substantial proportion of the variance for stride length and step length (17.6–28.8%) and contributed 10.6–14.7% to hoof height and 12.5% to forelimb and hindlimb walking speed, reflecting systematic developmental changes in these outcomes over time.

The Trial main effect contributed minimally across all outcomes, generally accounting for less than 1% of the total variance. Likewise, the Lamb × Trial and Month × Trial interactions were consistently small, typically contributing less than 2%, indicating minimal trial-to-trial measurement variability.

Residual variance varied considerably among outcome measures. Temporal variables and stride-related parameters demonstrated relatively low residual variance (approximately 8.8–32.1%), whereas hoof height and step width exhibited larger residual components. Residual variance was greatest for left and right forelimb step width (60.6% and 59.4%, respectively), followed by left hind hoof height (44.7%), suggesting greater unexplained variability or measurement error for these outcomes.

Overall, the G-study demonstrated that biological differences among lambs and their individual developmental trajectories were the predominant sources of variation, whereas trial-related sources of measurement error contributed minimally to the observed gait measurements.

#### B.2 Decision Study

The D-study evaluated the number of gait cycles required to achieve acceptable dependability ($\Phi \geq 0.80$) for each gait outcome (Table S2). Twenty-six of the 38 outcomes (68.4%) reached the reliability criterion using only one or two gait cycles, 32 outcomes (84.2%) required no more than three gait cycles, and 34 outcomes (89.5%) achieved acceptable dependability within five gait cycles. This indicates that reliable measurement can be obtained with substantially fewer gait cycles than the 10 collected in the present study.

The gait acquisition protocol employed in the present study, consisting of 10 gait cycles per assessment, demonstrated high dependability for most gait outcome measures (Table S2; $\Phi_{10}$). Of the 38 variables evaluated, 36 (94.7%) achieved acceptable reliability ($\Phi_{10} \geq 0.80$), 24 (63.2%) demonstrated excellent reliability ($\Phi_{10} \geq 0.90$), and 17 (44.7%) achieved near-perfect reliability ($\Phi_{10} \geq 0.95$). Only right hind hoof height ($\Phi_{10} = 0.77$) and left hip drop ($\Phi_{10} = 0.72$) remained below the acceptable reliability threshold despite averaging ten gait cycles.

**Table S1: Percentage contribution of each variance component to the total variance for all gait outcome measures estimated using Generalizability Theory.** Variance was partitioned into components attributable to Lamb, Month, Trial, the Lamb × Month, Lamb × Trial, and Month × Trial interactions, and Residual error. Percentages represent the proportion of the total variance explained by each source for each gait outcome.

| | Percentage of Variance (%) | | | | | | |
|---|---|---|---|---|---|---|---|
| **Outcome** | **Lamb** | **Month** | **Trial** | **Lamb × Month** | **Lamb × Trial** | **Month × Trial** | **Residual** |
| **Hoof Height** | | | | | | | |
| LF Hoof Height GT | 42.6 | 11.7 | 0.0 | 26.1 | 1.6 | 0.0 | 17.9 |
| RF Hoof Height GT | 52.6 | 14.7 | 0.0 | 20.7 | 0.4 | 0.0 | 11.7 |
| LH Hoof Height GT | 25.2 | 4.8 | 0.0 | 24.4 | 0.8 | 0.0 | 44.7 |
| RH Hoof Height GT | 9.9 | 10.6 | 0.7 | 50.4 | 0.8 | 0.0 | 27.6 |
| **Temporal Parameters** | | | | | | | |
| LF Cycle | 58.6 | 0.0 | 0.6 | 17.5 | 0.0 | 0.0 | 23.3 |
| RF Cycle | 58.3 | 0.0 | 0.8 | 17.7 | 0.0 | 0.0 | 23.3 |
| LH Cycle | 58.0 | 0.0 | 0.9 | 17.7 | 0.0 | 0.0 | 23.3 |
| RH Cycle | 58.7 | 0.0 | 0.6 | 16.4 | 0.0 | 0.2 | 24.1 |
| LF Stance Time | 58.4 | 0.0 | 0.6 | 17.0 | 0.0 | 0.1 | 23.9 |
| RF Stance Time | 57.0 | 0.0 | 0.5 | 18.4 | 0.0 | 0.0 | 24.0 |
| LH Stance Time | 55.1 | 0.0 | 0.5 | 21.7 | 0.0 | 0.3 | 22.4 |
| RH Stance Time | 60.5 | 0.0 | 0.4 | 17.2 | 0.3 | 0.0 | 21.6 |
| LF Swing Time | 47.2 | 0.0 | 0.4 | 20.3 | 0.0 | 0.0 | 32.1 |
| RF Swing Time | 52.2 | 0.0 | 0.7 | 15.3 | 0.0 | 0.0 | 31.8 |
| LH Swing Time | 40.6 | 0.5 | 1.0 | 20.2 | 0.6 | 0.0 | 37.0 |
| RH Swing Time | 43.1 | 0.8 | 1.0 | 20.2 | 0.0 | 0.0 | 34.9 |
| LF Step Time | 56.2 | 0.3 | 0.4 | 16.1 | 0.0 | 0.0 | 26.9 |
| RF Step Time | 52.5 | 0.0 | 0.6 | 19.7 | 0.3 | 0.0 | 27.0 |
| LH Step Time | 60.2 | 0.0 | 0.6 | 13.5 | 0.0 | 0.2 | 25.5 |
| RH Step Time | 51.3 | 0.0 | 0.6 | 19.6 | 0.2 | 0.0 | 28.4 |
| **Spatial Parameters** | | | | | | | |
| LF Stride Length | 55.3 | 26.8 | 0.0 | 8.6 | 0.2 | 0.0 | 9.1 |
| RF Stride Length | 55.0 | 27.0 | 0.1 | 8.6 | 0.5 | 0.0 | 8.8 |
| LH Stride Length | 53.3 | 26.5 | 0.0 | 8.5 | 0.5 | 0.0 | 11.2 |
| RH Stride Length | 53.4 | 26.1 | 0.0 | 9.0 | 1.0 | 0.0 | 10.4 |
| LF Step Length | 58.6 | 17.6 | 0.0 | 8.4 | 0.0 | 0.0 | 15.4 |
| RF Step Length | 44.3 | 28.8 | 0.1 | 11.0 | 1.1 | 0.0 | 14.8 |
| LH Step Length | 54.5 | 18.0 | 0.0 | 7.3 | 0.9 | 0.8 | 18.5 |
| RH Step Length | 51.5 | 20.4 | 0.0 | 9.8 | 0.3 | 0.6 | 17.4 |
| LF Step Width | 33.3 | 4.6 | 0.4 | 1.2 | 0.0 | 0.0 | 60.6 |
| RF Step Width | 35.0 | 3.4 | 0.0 | 2.1 | 0.0 | 0.0 | 59.4 |
| LH Step Width | 63.4 | 1.2 | 0.3 | 7.8 | 0.0 | 0.0 | 27.3 |
| RH Step Width | 66.6 | 0.8 | 0.4 | 5.6 | 0.1 | 0.0 | 26.6 |
| **Pelvic Kinematics** | | | | | | | |
| L Hip Drop GT | 7.3 | 0.0 | 0.4 | 63.4 | 0.0 | 0.4 | 28.5 |
| R Hip Drop GT | 18.6 | 0.0 | 0.1 | 60.1 | 0.4 | 0.0 | 20.9 |
| **Whole-Animal Outcomes** | | | | | | | |
| F Speed GT | 40.6 | 12.5 | 0.7 | 23.3 | 0.0 | 0.0 | 22.9 |
| H Speed GT | 40.9 | 12.5 | 0.7 | 23.5 | 0.0 | 0.0 | 22.4 |
| F DL Time GT | 60.3 | 0.0 | 0.6 | 18.4 | 0.0 | 0.0 | 20.7 |
| H DL Time GT | 57.1 | 0.0 | 0.2 | 27.7 | 0.6 | 0.0 | 14.3 |

*Abbreviations*: L = left; R = right; F = forelimb; H = hindlimb; DL = double-limb; GT = gait termination. Combined abbreviations denote limb laterality (e.g., LF = left forelimb, RF = right forelimb, LH = left hindlimb, and RH = right hindlimb).

**Table S2: Dependability coefficients ($\boldsymbol{\Phi}$) estimated from the Decision (D) study for protocols averaging one to ten gait cycles ($\boldsymbol{\Phi_1}$–$\boldsymbol{\Phi_{10}}$).** The current study protocol corresponds to $\boldsymbol{\Phi_{10}}$, as ten gait cycles were collected per assessment. The Critical Number of Trials ($\boldsymbol{nT}_{\text{Critical}}$) represents the minimum number of gait cycles required to achieve acceptable reliability ($\boldsymbol{\Phi} \geq \mathbf{0.80}$).

| Outcome | $\Phi_1$ | $\Phi_2$ | $\Phi_3$ | $\Phi_4$ | $\Phi_5$ | $\Phi_6$ | $\Phi_7$ | $\Phi_8$ | $\Phi_9$ | $\Phi_{10}$ | $n_T$ |
|---|---|---|---|---|---|---|---|---|---|---|---|
| **Hoof Height** | | | | | | | | | | | |
| LF Hoof Height GT | 0.69 | 0.81 | 0.87 | 0.9 | 0.92 | 0.93 | 0.94 | 0.95 | 0.95 | 0.96 | 2 |
| RF Hoof Height GT | 0.81 | 0.9 | 0.93 | 0.95 | 0.96 | 0.96 | 0.97 | 0.97 | 0.98 | 0.98 | 1 |
| LH Hoof Height GT | 0.36 | 0.53 | 0.62 | 0.69 | 0.73 | 0.77 | 0.79 | 0.82 | 0.83 | 0.85 | 8 |
| RH Hoof Height GT | 0.25 | 0.41 | 0.51 | 0.58 | 0.63 | 0.67 | 0.7 | 0.73 | 0.75 | 0.77 | 12 |
| **Temporal Parameters** | | | | | | | | | | | |
| LF Cycle | 0.71 | 0.83 | 0.88 | 0.91 | 0.92 | 0.94 | 0.94 | 0.95 | 0.96 | 0.96 | 2 |
| RF Cycle | 0.71 | 0.83 | 0.88 | 0.91 | 0.92 | 0.94 | 0.94 | 0.95 | 0.96 | 0.96 | 2 |
| LH Cycle | 0.71 | 0.83 | 0.88 | 0.91 | 0.92 | 0.93 | 0.94 | 0.95 | 0.96 | 0.96 | 2 |
| RH Cycle | 0.7 | 0.83 | 0.88 | 0.9 | 0.92 | 0.93 | 0.94 | 0.95 | 0.96 | 0.96 | 2 |
| LF Stance Time | 0.7 | 0.83 | 0.88 | 0.91 | 0.92 | 0.93 | 0.94 | 0.95 | 0.96 | 0.96 | 2 |
| RF Stance Time | 0.7 | 0.82 | 0.87 | 0.9 | 0.92 | 0.93 | 0.94 | 0.95 | 0.95 | 0.96 | 2 |
| LH Stance Time | 0.71 | 0.83 | 0.88 | 0.91 | 0.92 | 0.94 | 0.94 | 0.95 | 0.96 | 0.96 | 2 |
| RH Stance Time | 0.73 | 0.84 | 0.89 | 0.92 | 0.93 | 0.94 | 0.95 | 0.96 | 0.96 | 0.96 | 2 |
| LF Swing Time | 0.59 | 0.74 | 0.81 | 0.85 | 0.88 | 0.9 | 0.91 | 0.92 | 0.93 | 0.94 | 3 |
| RF Swing Time | 0.62 | 0.76 | 0.83 | 0.87 | 0.89 | 0.91 | 0.92 | 0.93 | 0.94 | 0.94 | 3 |
| LH Swing Time | 0.51 | 0.68 | 0.76 | 0.81 | 0.84 | 0.86 | 0.88 | 0.89 | 0.9 | 0.91 | 4 |
| RH Swing Time | 0.55 | 0.71 | 0.78 | 0.83 | 0.86 | 0.88 | 0.89 | 0.91 | 0.92 | 0.92 | 4 |
| LF Step Time | 0.67 | 0.8 | 0.86 | 0.89 | 0.91 | 0.92 | 0.93 | 0.94 | 0.95 | 0.95 | 2 |
| RF Step Time | 0.65 | 0.79 | 0.85 | 0.88 | 0.9 | 0.92 | 0.93 | 0.94 | 0.94 | 0.95 | 3 |
| LH Step Time | 0.7 | 0.82 | 0.87 | 0.9 | 0.92 | 0.93 | 0.94 | 0.95 | 0.95 | 0.96 | 2 |
| RH Step Time | 0.64 | 0.78 | 0.84 | 0.88 | 0.9 | 0.91 | 0.92 | 0.93 | 0.94 | 0.95 | 3 |
| **Spatial Parameters** | | | | | | | | | | | |
| LF Stride Length | 0.86 | 0.92 | 0.95 | 0.96 | 0.97 | 0.97 | 0.98 | 0.98 | 0.98 | 0.98 | 1 |
| RF Stride Length | 0.85 | 0.92 | 0.95 | 0.96 | 0.97 | 0.97 | 0.98 | 0.98 | 0.98 | 0.98 | 1 |
| LH Stride Length | 0.82 | 0.9 | 0.93 | 0.95 | 0.96 | 0.96 | 0.97 | 0.97 | 0.98 | 0.98 | 1 |
| RH Stride Length | 0.82 | 0.9 | 0.93 | 0.95 | 0.96 | 0.97 | 0.97 | 0.97 | 0.98 | 0.98 | 1 |
| LF Step Length | 0.79 | 0.88 | 0.92 | 0.94 | 0.95 | 0.96 | 0.96 | 0.97 | 0.97 | 0.97 | 2 |
| RF Step Length | 0.74 | 0.85 | 0.89 | 0.92 | 0.93 | 0.94 | 0.95 | 0.96 | 0.96 | 0.97 | 2 |
| LH Step Length | 0.74 | 0.85 | 0.89 | 0.92 | 0.93 | 0.94 | 0.95 | 0.96 | 0.96 | 0.97 | 2 |
| RH Step Length | 0.74 | 0.85 | 0.9 | 0.92 | 0.94 | 0.95 | 0.95 | 0.96 | 0.96 | 0.97 | 2 |
| LF Step Width | 0.35 | 0.52 | 0.62 | 0.69 | 0.73 | 0.77 | 0.79 | 0.81 | 0.83 | 0.85 | 8 |
| RF Step Width | 0.37 | 0.54 | 0.64 | 0.7 | 0.75 | 0.78 | 0.8 | 0.82 | 0.84 | 0.85 | 7 |
| LH Step Width | 0.7 | 0.82 | 0.87 | 0.9 | 0.92 | 0.93 | 0.94 | 0.95 | 0.95 | 0.96 | 2 |
| RH Step Width | 0.71 | 0.83 | 0.88 | 0.91 | 0.92 | 0.94 | 0.95 | 0.95 | 0.96 | 0.96 | 2 |
| **Pelvic Kinematics** | | | | | | | | | | | |
| L Hip Drop GT | 0.2 | 0.34 | 0.43 | 0.5 | 0.56 | 0.6 | 0.64 | 0.67 | 0.7 | 0.72 | 16 |
| R Hip Drop GT | 0.47 | 0.64 | 0.72 | 0.78 | 0.81 | 0.84 | 0.86 | 0.87 | 0.89 | 0.9 | 5 |
| **Whole-Animal Outcomes** | | | | | | | | | | | |
| F Speed GT | 0.63 | 0.77 | 0.84 | 0.87 | 0.9 | 0.91 | 0.92 | 0.93 | 0.94 | 0.94 | 3 |
| H Speed GT | 0.64 | 0.78 | 0.84 | 0.88 | 0.9 | 0.91 | 0.93 | 0.93 | 0.94 | 0.95 | 3 |
| F DL Time GT | 0.74 | 0.85 | 0.89 | 0.92 | 0.93 | 0.94 | 0.95 | 0.96 | 0.96 | 0.97 | 2 |
| H DL Time GT | 0.79 | 0.88 | 0.92 | 0.94 | 0.95 | 0.96 | 0.96 | 0.97 | 0.97 | 0.97 | 2 |

$\mathbf{\Phi_{10}}$ represents the dependability coefficient for the gait acquisition protocol implemented in the present study (10 gait cycles per assessment).

$\mathbf{n_T}$ denotes the minimum number of gait cycles required to achieve a dependability coefficient of at least 0.80.

*<u>Abbreviations:</u>* L = left; R = right; F = forelimb; H = hindlimb; DL = double-limb; GT = gait termination. Combined abbreviations denote limb laterality (e.g., LF = left forelimb, RF = right forelimb, LH = left hindlimb, and RH = right hindlimb).
