## Supplement 2 - LMM Equation for "Quantitative Gait Analysis in a Sheep Model of In-Utero Spina Bifida Repair"

**Supplementary Material 2.**

The linear mixed model equation is expressed as:

$$\mathrm{Y}_{gait} = \beta_0 + \beta_1(\mathrm{Group}) + \beta_2(\mathrm{Month}) + \beta_3(\mathrm{Group} * \mathrm{Month}) + \upsilon + \varepsilon \qquad \text{Equation S1}$$

$Y_{\text{gait}}$ represents the gait outcome (e.g., gait speed, step length) measured monthly for each sheep; $\beta_0$ is the expected gait outcome for the reference group at the reference month; Group is a categorical factor (control vs. treatment); Month is a categorical factor (3-6 months); Group*Month interaction captures whether the longitudinal change differs between groups. $\upsilon$ is the random interception for variability between sheep, and $\varepsilon$ is the residual error. Correlation among repeated measurements within each sheep was modeled using AR (1) covariance structure.
