## Supplement 3 - Mahalanobis Distance for "Quantitative Gait Analysis in a Sheep Model of In-Utero Spina Bifida Repair"

**Supplementary Material 3.**

**Table S3. Descriptive statistics of left-limb, right-limb, and bilateral Mahalanobis distances for each treated sheep.**

| Sheep # | Mahalanobis distance | | |
|---|---|---|---|
| | Left limb | Right limb | Bilateral limbs |
| 1 | 2.1 ± 0.9 | 1.8 ± 0.6 | 3.9 ± 1.1 |
| 2 | 6.6 ± 4.2 | 3.2 ± 1.8 | 14.4 ± 9.8 |
| 3 | 1.5 ± 0.4 | 1.3 ± 0.5 | 3.0 ± 0.8 |
| 4 | 1.3 ± 0.5 | 2.0 ± 0.6 | 3.6 ± 1.0 |
| 5 | 1.3 ± 0.5 | 1.9 ± 0.9 | 3.9 ± 1.4 |
| 6 | 2.2 ± 0.9 | 2.0 ± 0.7 | 3.6 ± 1.2 |
| 7 | 2.2 ± 0.5 | 2.2 ± 0.8 | 3.9 ± 0.8 |
| 8 | 1.8 ± 0.7 | 2.1 ± 1.1 | 3.5 ± 1.7 |
| 9 | 2.0 ± 0.9 | 1.7 ± 0.4 | 3.2 ± 0.7 |
| 10 | 1.3 ± 0.4 | 1.5 ± 0.5 | 2.3 ± 0.5 |
| 11 | 1.5 ± 0.4 | 3.3 ± 2.9 | 4.6 ± 3.2 |
| 12 | 1.6 ± 0.6 | 1.3 ± 0.4 | 2.5 ± 0.8 |
| 13 | 1.5 ± 0.6 | 1.3 ± 0.3 | 2.6 ± 0.7 |
| 14 | 1.2 ± 0.3 | 1.3 ± 0.4 | 2.3 ± 0.5 |

Data are presented as mean ± standard deviation. Sheep #2 was euthanized before 5 months because of disease progression.
